# Cardiac *GLP1R* gene Expression: A Cross-Species Single-Cell Transcriptomic Analysis

**DOI:** 10.64898/2026.01.18.700176

**Authors:** Christopher Dostal, Matthias Ernst, Johanna Reiner, Ryan Snelling, Chenming Hu, Peter Pokreisz, Bruno K. Podesser, Attila Kiss

## Abstract

Glucagon-like peptide-1 receptor (GLP1R) agonists improve glycaemic control, induce weight loss, and consistently reduce major adverse cardiovascular events. However, the mechanistic basis of their cardioprotective effects remains incompletely understood, particularly whether benefits arise solely from systemic actions or also involve direct cardiac GLP1R signalling. To address this, we performed integrated single-cell and single-nucleus transcriptomics to map *GLP1R* gene expression across human and murine organs, cardiac cell types, disease states, and hiPSC-derived cardiac organoids. In humans, *GLP1R* expression was predominantly pancreatic, with low cardiac expression largely restricted to cardiomyocytes and consistently upregulated across ischaemic, dilated, and hypertrophic cardiomyopathy. In contrast, murine cardiac *Glp1r* expression was confined to endocardial cells and remained unchanged in heart disease. Other cardiac cell types, including fibroblasts, endothelial cells, and mural cells, showed minimal *GLP1R* expression in both species. Human cardiac organoids recapitulated ventricular GLP1R patterns closer to adult human myocardium than murine tissue. Together, these findings indicate that GLP1R is primarily extracardiac but selectively induced in failing human myocardium, supporting a model in which myocardial GLP1R signalling augments systemic mechanisms to confer GLP1R agonist-mediated cardioprotection.

## Introduction

Recent advances in pharmacotherapy with sodium-glucose cotransporter 2 inhibitors (SGLT2i) and glucagon-like peptide-1 receptor (GLP1R) agonists have reshaped the management of cardiometabolic disease, a major driver of heart failure with both preserved (HFpEF) and reduced (HFrEF) ejection fraction^1^. The cardiovascular benefits of SGLT2i are well established, with consistent improvements in heart failure outcomes across diverse patient populations^2–4^. In contrast, GLP1R agonists are primarily indicated for glycemic control and weight reduction; however, accumulating evidence supports a clinically meaningful cardiovascular benefit^5,6^, particularly among patients with obesity at high cardiovascular risk^6–10^. Data from cardiovascular outcome trials suggest a more pronounced benefit in HFpEF compared with HFrEF^5,11–15^. Despite these advances, the underlying mechanisms of GLP1R agonism remain incompletely understood, in part because canonical signaling pathways in the heart have not been definitively characterized^16^.

In preclinical models, GLP1R agonists similarly confer robust protection across diverse forms of cardiac injury, including metabolic dysfunction, myocardial ischemia, pressure overload, and doxorubicin cardiotoxicity^17–23^. Proposed mechanisms include improvements in myocardial metabolism, microvascular function, and inflammatory pathways^21–25^. Notably, important species differences exist: in mice, Glp1r expression is predominantly localized to endocardial and endothelial cells^21,26^, whereas in humans, myocardial GLP1R expression is sparse, heterogeneous, and inconsistently detected^27,28^.

It therefore remains unclear whether GLP1R agonists act directly on the myocardium or mainly through systemic mechanisms, reflecting incomplete data on species- and cell-type-specific expression, as well as limited insight into expression changes in failing hearts. To address these gaps, we integrated single-cell and single-nucleus transcriptomic datasets to systematically map GLP1R mRNA expression across species, developmental stages, disease states and cardiac model systems, providing a framework for evaluating direct cardiac GLP1R signalling.

## Methods

### Single-cell and single-nucleus transcriptomic data analysis

Publicly available single-cell and single-nucleus RNA sequencing datasets from human and mouse hearts were analysed to characterize tissue- and cell-type–specific expression of the glucagon-like peptide-1 receptor (GLP1R/Glp1r). Human and mouse were selected to combine clinical relevance with cross-species validation in established preclinical heart failure models. Datasets spanned physiological and pathological conditions, including non-failing hearts, dilated and hypertrophic cardiomyopathy, ischemic cardiomyopathy, mouse models of HFpEF and HFrEF, and human iPSC-derived cardiac organoids. An overview of all datasets, including species, tissue sources and sample numbers, is provided in Table 1.

**Table 1.** Overview of included datasets.

| Species | Description | Sex | Tissue | Cells/nuclei | n | Reference |
| --- | --- | --- | --- | --- | --- | --- |
| Human | Reference human cell atlas | Mixed | 28 organs | 1.1M | 24 | Tabula Sapiens Consortium <sup>29</sup> |
| Mouse | Reference mouse cell atlas | Mixed | 20 organs | 100,605 | 8 | Tabula Muris Consortium <sup>30</sup> |
| Human | ICM | Mixed | LV | 99,684 | 7 ICM<br>8 NF | Simonson et al. <sup>31</sup> |
| Human | HCM and DCM | Mixed | LV | 592,689 | 11 DCM<br>15 HCM<br>15 NF | Chaffin et al. <sup>32</sup> |
| Mouse | Mouse models of HFpEF and HFrEF | Male | LV | 44,133 | 2 WT Sham<br>2 WT TAC<br>2 ob/ob Sham<br>2 ob/ob TAC | Gao et al. <sup>33</sup> |
| Human | iPSC-derived cardiac organoid | WTC-11 (male) iPSC cell line | Fetal-like cardioid | 36,000 - 146,000 | 2 (16-36 cardioids each) | Schmidt et al. <sup>34</sup> |

n: number of biological replicates; HCM: hypertrophic cardiomyopathy; DCM: dilated cardiomyopathy; LV: left ventricle; NF: non failing; ICM: Ischemic cardiomyopathy; iPSC: induced pluripotent stem cell.

### Datasets

Six datasets were included (Table 1):

### Tabula Sapiens and Tabula Muris datasets

Multi-organ single-cell reference atlases from human (Tabula Sapiens) and mouse (Tabula Muris) were obtained from respective public repositories. Heart, lung, pancreas and other relevant tissues were extracted to compare organ-level enrichment of GLP1R/Glp1r expression across species.

### Human dilated, hypertrophic, and ischemic cardiomyopathy datasets

Single-nucleus RNA-seq datasets of non-failing, dilated and hypertrophic cardiomyopathy hearts, as well as ischemic cardiomyopathy and matched non-failing donor hearts, were obtained from CELLxGENE and GEO as indicated in Table 1. These datasets were used to assess GLP1R expression across major cardiac cell types and to compare disease versus control hearts.

### Mouse heart failure models

Single-nucleus RNA-seq data from left ventricular tissue of HFpEF and HFrEF mouse models (WT and obese (ob/ob) mice subjected to sham or transverse aortic constriction (TAC) surgery) were obtained from the GEO repository (GSE236585). These data were used to examine Glp1r expression in cardiomyocytes and non-myocyte populations under experimental pressure overload and metabolic stress.

### Human iPSC-derived cardiac organoids

Single-cell RNA-seq data from fetal-like human iPSC-derived cardiac organoids (cardioids) generated from the WTC-11 iPSC line were obtained from the GEO repository (GSE239890)^34^. These data were analysed to determine GLP1R expression in developing human cardiac-like tissues.

### Data processing and cell-type annotation

All analyses were performed in Python (v3.10.19) using Scanpy (v1.11.5), and standard single-cell workflows. Human age-stratified cardiac expression was evaluated in the non-failing cohort of Chaffin et al., comprising 4 donors <50 years and 11 donors >50 years (33–72 years). Disease-associated expression was assessed in independent datasets of ICM vs non-failing, and in an integrated NF vs DCM vs HCM dataset, as well as in a mouse TAC model (WT TAC vs ob/ob TAC). Developmental cardiac expression was analyzed in cardiac organoids, using the published subtype annotations (LV, RV, atria, OFT, AVC).

### Quantification of GLP1R expression and comparative analyses

For each dataset, GLP1R (human) or Glp1r (mouse) expression was summarised per major cardiac cell type as the proportion of GLP1R-positive cells and mean scaled expression and visualised using dot plots. In Tabula Sapiens/Muris and organoid datasets these metrics were compared descriptively across organs and cell types, whereas in disease datasets we additionally performed cell type–specific differential expression between non-failing and disease groups (Wilcoxon rank-sum on log-normalized data) to obtain fold-changes and P values, interpreting GLP1R mainly in terms of direction and magnitude of change and positive-cell frequency given its sparse expression.

## Results

GLP1R expression was assessed systemically in healthy human and mouse reference cohorts, followed by heart-specific profiling of cell types across age groups (Fig. 1). In both species, GLP1R expression was most prominent in the pancreas and lung, whereas the heart and other tissues exhibited comparatively low transcript abundance. In the Tabula Sapiens atlas, GLP1R expression was enriched in pancreatic delta and beta cells, with additional expression detected in bronchial smooth muscle cells and selected resident lymphocytes of the liver, kidney, and vasculature (Fig. 1A). In the Tabula Muris atlas, Glp1r expression showed a comparable distribution, with highest levels in pancreatic and lung endothelial cell populations (Fig. 1B). GLP1R expression in the human heart was sparse and primarily detected in cardiomyocytes (CM) (Fig. 1C). In contrast, murine cardiac Glp1r expression was similarly low but confined to endocardial cells (Fig. 1E).

**Figure 1:**
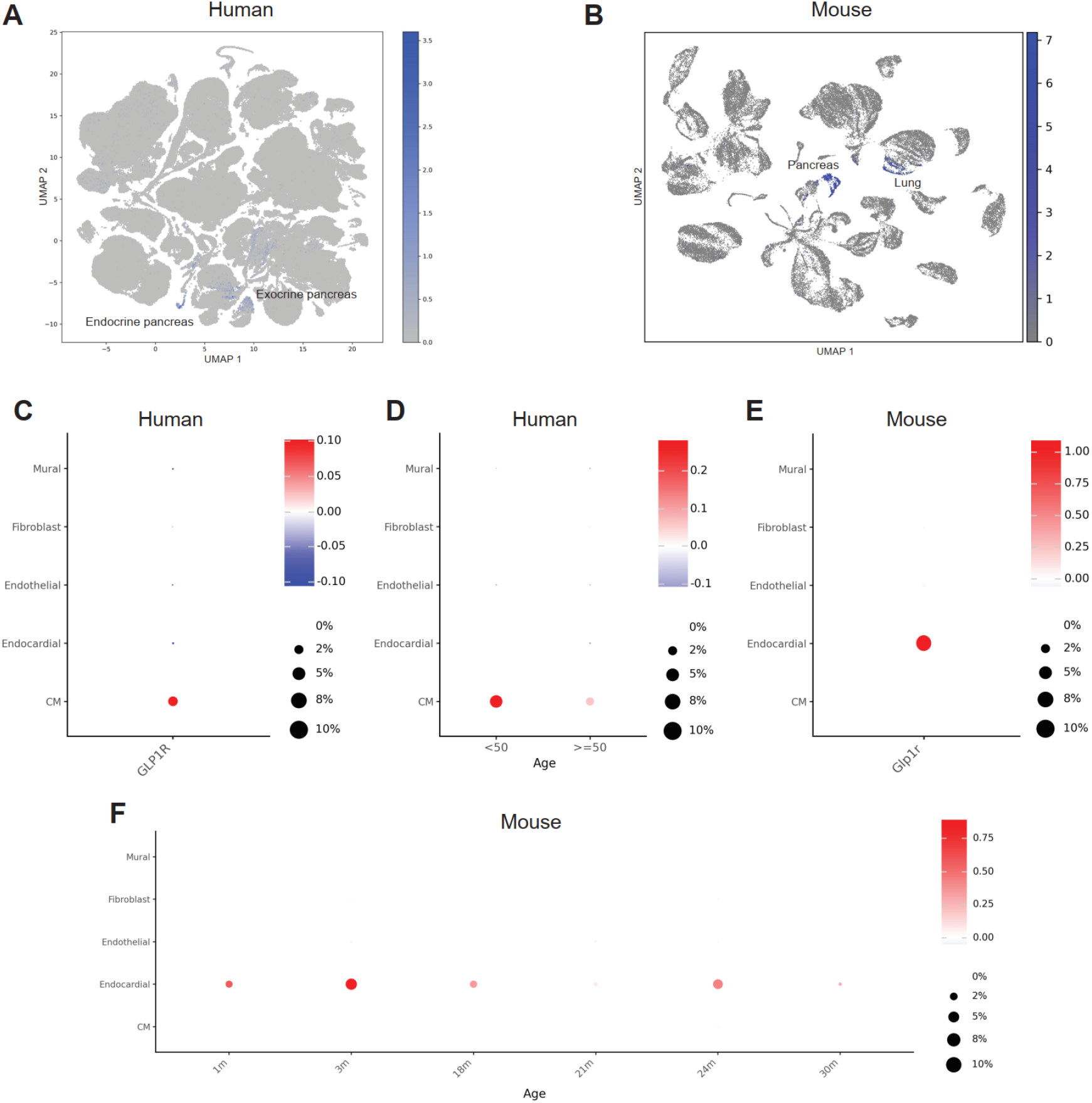
GLP1R expression in the healthy heart across species, tissue, and age. A: UMAP hex-bin projection of GLP1R expression between different organs of human (n = 24; Tabula Sapiens) and (B) UMAP projection of GLP1R expression between different organs of mouse (n = 8; Tabula Muris). C/E: Feature plots quantifying cardiac GLP1R expression in human (n =15; Chaffin et al.)) (C) and mouse (n = 8; Tabula Muris) (E). Cell types were chosen based on relative abundance, expression level, and physiological relevance. D/F: Feature plots of age-dependent GLP1R expression in cardiac cell types in human (n =15; Chaffin et al.) (D) and mouse (n = 8; Tabula Muris) (F).

We next evaluated age-specific patterns of GLP1R expression using the non-failing human heart cohort from Chaffin et al. ^32^, comprising four donors <50 years, and 11 donors >50 years (range: 33-72 years). GLP1R expression was restricted to cardiomyocytes and remained largely stable across age, with an apparent increase driven by a single donor in the younger cohort (Fig. 1D). In the Tabula Muris dataset, endocardial Glp1r expression was highest during the first 12 weeks of life followed by a decline and a modest secondary increase at 24 months (Fig. 1F).

Together, these data indicate that GLP1R expression is predominantly confined to pancreatic and pulmonary cell populations across species, with only minimal expression in the heart. In humans, cardiac GLP1R levels remain low and largely stable throughout adulthood, whereas in mice, Glp1r is transiently enriched in endocardial cells during early postnatal life.

GLP1R expression was next examined across cardiomyopathy subtypes relative to the non-failing heart (Fig. 2). Cardiac expression was analyzed in human datasets of ischemic (ICM), dilated (DCM), and hypertrophic (HCM) cardiomyopathy, as well as in murine models of heart failure with reduced (WT TAC) and preserved (ob/ob TAC) LV ejection fraction ^31–33^. In ICM, expression was largely confined to cardiomyocytes with a modest but consistent increase compared with non-failing (NF) hearts ^31^. Non-myocyte populations, including fibroblasts, endothelial, and immune cells, exhibited minimal expression in both groups (Fig. 2A). In DCM and HCM populations, cardiomyocytes similarly displayed the highest GLP1R expression, with elevated levels in HCM relative to NF hearts (logFC of 0.61), and most pronounced expression in DCM (logFC over NF of 1.53) (Fig. 2B).

**Figure 2:**
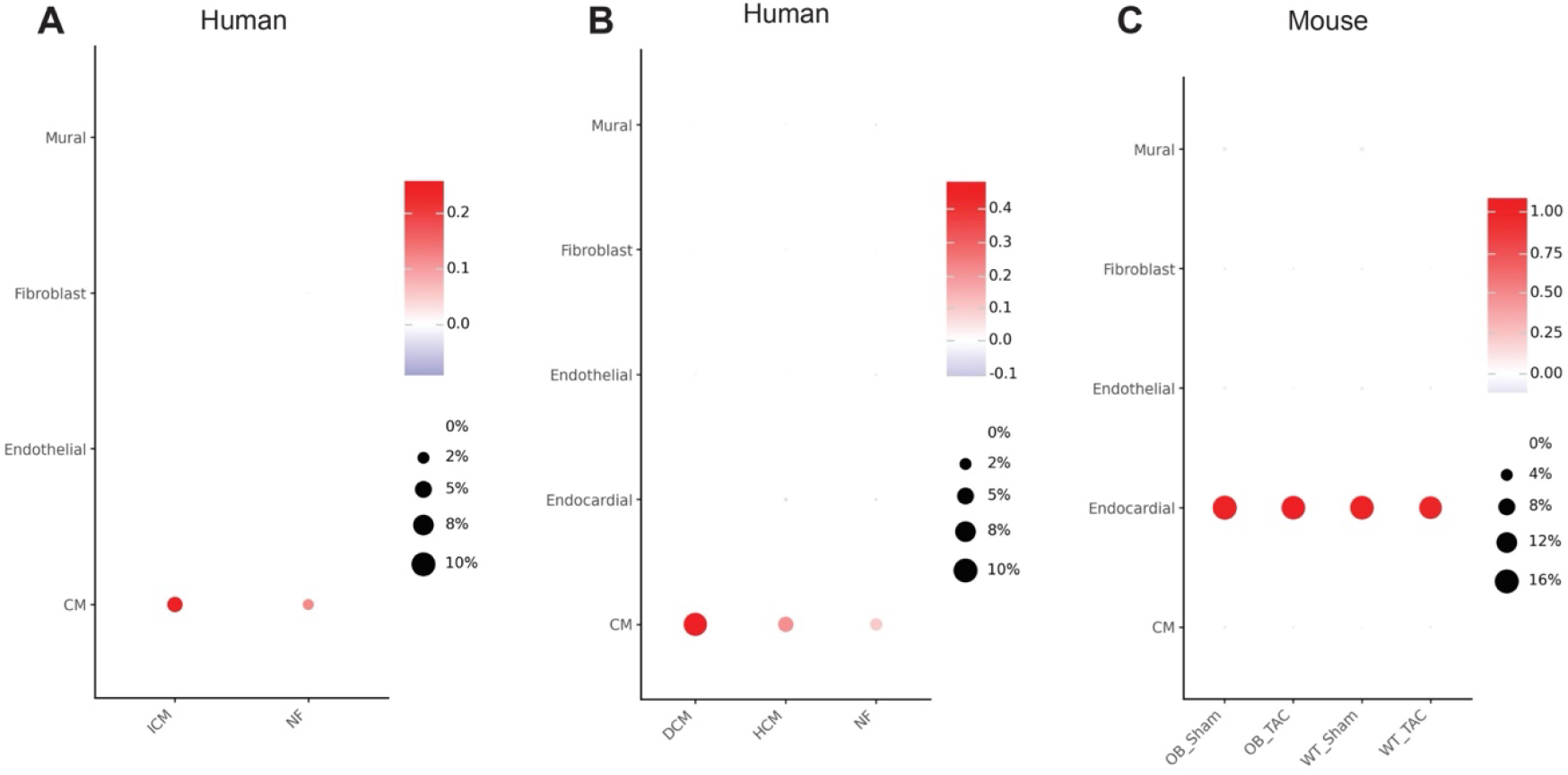
GLP1R expression in various heart diseases across species. A: Feature plot of GLP1R expression in human non-failing (NF) (n = 8; Simonson et al.) versus ischemic cardiomyopathy (ICM) (n = 7; Simonson et al.) hearts. B: Feature plot of GLP1R expression in human NF (n = 16; Chaffin et al.), hypertrophic cardiomyopathy (HCM) (n = 15; Chaffin et al.), and dilated cardiomyopathy (DCM) (n = 11; Chaffin et al.) hearts. **C:** Feature plot of GLP1R expression in mice with or without obesity (OB) and/or transverse aortic constriction (TAC) (n = 2 in each group; Gao et al.). OB + TAC and WT + TAC model HFpEF and HFrEF, respectively.

In mice, Glp1r expression was consistent across all experimental groups and remained confined to the endocardium, irrespective of HF status (Fig. 2C) ^33^. RNA sequencing was performed at 18 weeks of age, corresponding to the period of endocardial expression observed in the Tabula Muris dataset (Fig. 1F). Collectively, these findings indicate a modest disease-associated induction of GLP1R transcripts in human cardiomyocytes, contrasted with stable disease-independent endocardial expression in mice.

The observed differences in GLP1R expression between human and mouse hearts prompted evaluation of additional cardiac model systems. Schmidt et al. ^34^ developed human cardioids comprising five chamber-specific subtypes: left ventricle (LV), right ventricle (RV), atria, outflow tract (OFT), and atrioventricular tract (AVC) (Fig. 3A). Each subtype underwent single cell RNA sequencing at day 9.5 with two biological replicates (16-36 cardioids each), except the LV subtype, which was sequenced at day 7.5, and the atrial subtype, which included only a single replicate. GLP1R expression was highest in the contractile LV and RV, whereas atrial, OFT, and AVC subtypes exhibited minimal expression (Fig. 3B). Despite its fetal-like phenotype, this iPSC-derived human cardiac organoid model demonstrates GLP1R expression resembling that of the adult human heart.

**Figure 3:**
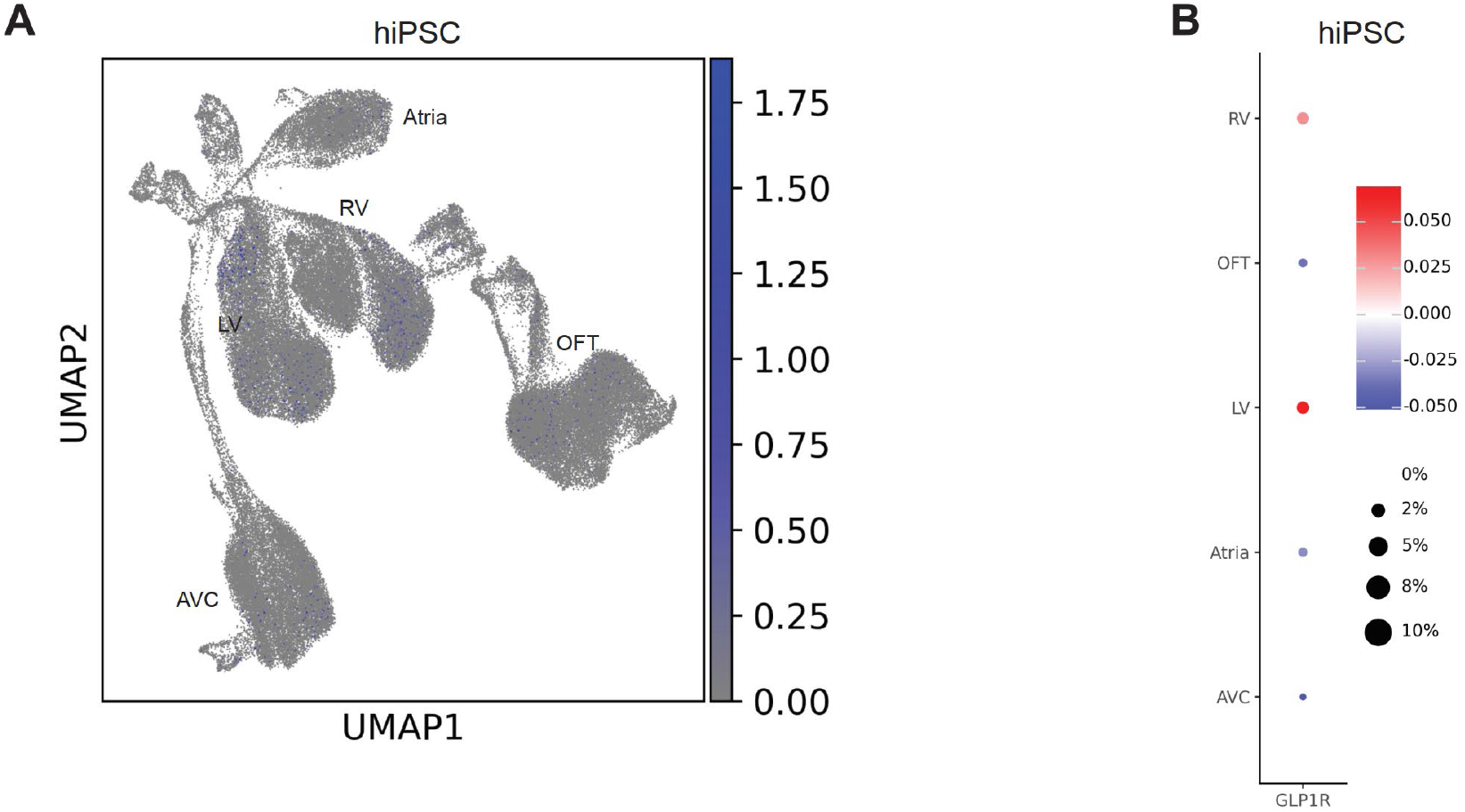
GLP1R expression in human stem cell-derived cardiac organoids. A: UMAP projection outlining cardiac organoid subtypes: LV, RV, Atrial, OFT, and AVC (Atrial n = 1; others n = 2 each; Schmidt et al.). B: Feature plot showing GLP1R expression across these subtypes. LV: left ventricle; RV: right ventricle; OFT: outflow tract; AVC: atrioventricular canal.

## Discussion

As GLP1R agonism emerges as a promising therapeutic strategy for HFpEF^35^, this cross-species analysis helps clarify the relative contributions of direct cardiac versus systemic mechanisms. We examined GLP1R expression across healthy and diseased human and mouse hearts, as well as human iPSC-derived cardiac organoids, using publicly available single-cell and single-nucleus transcriptomic datasets.

Two principal observations emerge. First, GLP1R expression in the heart is uniformly low and exhibits species-dependent distribution: mice show Glp1r transcripts enriched in endocardial-endothelial cells, whereas human datasets reveal sparse expression in cardiomyocytes. Second, analysis of human cardiomyopathy samples (ICM, DCM, HCM) reveals a modest, disease-associated upregulation of cardiomyocyte GLP1R transcripts, whereas murine HFpEF and HFrEF models show no analogous change, indicating important species-specific differences in cardiac GLP1R regulation.

Our results are consistent with prior studies demonstrating largely absent GLP1R expression in human coronary endothelial cells, vascular smooth muscle cells, and cardiac fibroblasts, with minimal but detectable transcripts in atrial and ventricular cardiomyocytes^27^. These expression patterns should be interpreted alongside the consistent clinical cardioprotective effects of GLP1R agonists, including reductions in major adverse cardiovascular events in diabetes (LEADER, SUSTAIN-6) ^7,8^ and symptomatic and functional improvement in obese HFpEF patients (STEP-HFpEF) ^5^. The observed induction of GLP1R expression in human cardiomyopathy raises the possibility that cardiac GLP1R signalling contributes to cardioprotection in addition to broader systemic effects. However, the cohorts analysed in this study largely represent patients with end-stage heart failure, with only limited representation of HFpEF - primarily among individuals with hypertrophic cardiomyopathy. Defining the precise role of canonical GLP1R signaling in HFpEF would require more comprehensive in well-phenotyped patient cohorts, incorporating analyses of both cardiac tissue and pericardial fat to fully capture the spectrum of cardiac and systemic mechanisms.

Human iPSC-derived cardiac organoids offer an alternative model in which GLP1R expression is more readily detected in ventricular cardiomyocytes. Interestingly, these cardioids reveal chamber-specific GLP1R expression, suggesting a potential role for GLP1R in cardiac development. However, organoids largely retain a fetal-like transcriptional and functional state and this model does not capture key HFpEF features such as ageing, female sex, or multimorbidity, limiting their utility for disease-specific inference. In this context, genetic models of cardiomyopathy, such as HCM and DCM, may provide a more physiologically relevant system to study disease-specific regulation and functional consequences of cardiac GLP1R. Our study may also highlight the limitations of rodent models in selected cardiovascular pathologies and underscores the importance of human-derived data, providing a framework for interpreting GLP1R agonist effects and motivating future studies that integrate protein-level analyses, spatial transcriptomics, and advanced human cardiac models.

In murine models, the cardioprotective effects of GLP1R agonists appear to depend on canonical signalling within endothelial or endocardial compartments ^21,36^, whereas cardiomyocyte-specific deletion of Glp1r does not attenuate therapeutic efficacy ^28^. This apparent discrepancy supports a model in which both cardiac and extra-cardiac mechanisms contribute to cardioprotection, while underscoring critical species-specific differences that may limit the translational relevance of mouse models for human GLP1R signaling. Importantly, these differences may arise not only from transcriptional regulation but also from post-translational processes - including receptor trafficking, internalization, and degradation that dynamically shape GLP1R abundance and functional responsiveness in cardiomyocytes and other cardiac cell types.

Our findings are subject to certain limitations that warrant consideration. All datasets rely on single-cell or single-nucleus transcriptomics, which are prone to dropout for low-abundance GPCRs ^37^. Protein-level validation is lacking, and antibody-based detection remains challenging due to known cross-reactivity issues ^38^.

Human datasets are biased toward end-stage HFrEF, restricting insight into earlier HFpEF phenotypes such as obesity-related HFpEF, where GLP1R agonists may be most effective. The end-stage sampling also precluded capturing the relatively rapid changes in transcript abundance and dynamics that occur during the compensatory phases of cardiac disease. In addition, the current analysis was restricted to a single murine model encompassing HFpEF and HFrEF, which may not fully reflect the biological heterogeneity of heart failure. As such, disease-specific differences in GLP1R expression or function in advanced heart failure cannot be excluded.

In summary, cardiac GLP1R expression is minimal, highly cell-type restricted, and strongly species dependent. In mice, GLP1R is largely confined to endocardial and endothelial cells, whereas in humans it is limited and predominantly localized to cardiomyocytes; in human cardiac organoids, expression is selectively enriched in ventricular cardiomyocytes. Failing human hearts exhibit modest disease-associated induction of GLP1R, in contrast to the stable expression observed across murine models of heart failure. Collectively, these data indicate that direct cardiac GLP1R signaling is limited to discrete receptor-positive populations and support the interpretation that indirect systemic mechanisms (cardiometabolic, vascular, renal, and anti-inflammatory effects) contribute substantially to the clinically observed cardioprotection^6,36,39,40^.

## Author Contributions

**C. D**, Conceived and designed research, Analyzed data, Interpreted results of experiments, Prepared figures, Drafted manuscript, Approved final version. **M.E**., Interpreted results of experiments, Drafted manuscript, Edited and revised manuscript, Approved final version. **J.R**., Analyzed data, Interpreted results of experiments, Prepared figures. **R.S**., Interpreted results of experiments, Drafted manuscript, Edited and revised manuscript, Prepared figures. **C.H**., Drafted manuscript, Edited and revised manuscript. **P.P**., Edited and revised manuscript. **B.K.P**., Edited and revised manuscript, Approved final version. **A.K**., Edited and revised manuscript, Approved final version.

## Data Availability

All datasets analysed in this study are publicly available from the cited repositories. Analysis scripts are available from the corresponding author upon reasonable request.

